# Exploring the role of task success in implicit motor adaptation

**DOI:** 10.1101/2023.02.01.526533

**Authors:** Naser Al-Fawakhiri, Ambri Ma, Jordan A. Taylor, Olivia A. Kim

**Affiliations:** Department of Psychology, Yale University, New Haven, CT 06511; Department of Psychology, Princeton University, Princeton, NJ 08544

**Author notes:** Correspondence to: Olivia Kim 428 Peretsman-Scully Hall Princeton, NJ 08502. **AUTHOR CONTRIBUTIONS**: Conceptualization, NA, JT, & OK; Methodology, NA, JT, & OK; Software, NA & OK; Validation, NA & OK; Formal Analysis, NA & OK; Investigation, NA, AM, & OK; Resources, JT; Data Curation, NA & OK; Supervision, JT & OK; Project Administration, JT & OK; Funding Acquisition, JT & OK; Writing, NA, AM, JT, & OK. **CONFLICT OF INTEREST**: The authors declare no competing financial interests.

**Keywords:** Sensorimotor adaptation, motor learning, reinforcement

## Abstract

While implicit motor adaptation is driven by sensory-prediction errors (SPEs), recent work has shown that task success modulates this process. Task success has typically been defined as hitting a target, which signifies the goal of the movement. Visuomotor adaptation tasks are uniquely situated to experimentally manipulate task success independently from SPE by changing the target size or the location of the target. These two, distinct manipulations may influence implicit motor adaptation in different ways, so, over four experiments, we sought to probe the efficacy of each manipulation. We found that changes in target size which caused the target to fully envelop the cursor only affected implicit adaptation for a narrow range of SPE sizes, while jumping the target to overlap with the cursor more reliably and robustly affected implicit adaptation. Taken together, our data indicate that, while task success exerts a small effect on implicit adaptation, these effects are susceptible to methodological variations. Future investigations of the effect of task success on implicit adaptation could benefit from employing target jump manipulations instead of target size manipulations.

## NEW & NOTEWORTHY

Recent work has suggested that task success, namely, hitting a target, influences implicit motor adaptation. Here, we observed that implicit adaptation is modulated by target jump manipulations, where the target abruptly “jumps” to meet the cursor; however, implicit adaptation was only weakly modulated by target size manipulations, where a static target either envelops or excludes the cursor. We discuss how these manipulations may exert their effects through different mechanisms.

## INTRODUCTION

The acquisition and maintenance of motor skills requires learning to reduce motor errors (1-4). However, motor learning draws from multiple kinds of errors, including sensory-prediction errors (SPE) and task errors (TE) (5). SPEs represent the discrepancy between the observed and the intended movement outcome. TEs indicate whether the movement achieved a higher-level goal, such as hitting a target. Classically, sensorimotor adaptation has been conceived of as two processes, with a fast, explicit re-aiming process that reduces TE and a slow, implicit recalibration process that reduces SPE (6-8).

However, recent work suggests that the implicit process also responds to TEs (9-14). Implicit adaptation can be isolated using an “error-clamp,” in which the observed movement trajectory is fixed at a constant degree of angular error from the center of a reaching target (15). Kim and colleagues (11) combined the error-clamp approach alongside manipulations of Target Size to show implicit adaptation can be suppressed by task success signals. In one condition, the target was small, so the error-clamp caused the cursor to miss the target. In another condition, the target was larger, and the error-clamped cursor feedback was fully enclosed in the target. Participants in the latter condition exhibited less implicit adaptation than the former, suggesting that TEs affect implicit adaptation. Task success has also been manipulated using the “Target Jump” paradigm, where the location of the target changes during the reach to intersect the cursor (9, 13). When the target “jumps” such that the cursor hits the target, implicit adaptation is suppressed, further supporting the claim that implicit adaptation is sensitive to TEs. Notably, target jumps alone, in the absence of SPE, are not sufficient to drive implicit adaptation (13, 16)

Previous reports using Target Size and Target Jump perturbations both modulate implicit adaptation through perceptions of TE or task success (9, 11), with adaptation being suppressed upon successful movements. However, it is apparent that Target Size and Target Jump manipulations are distinct and may have different effects on implicit adaptation. If task success influences implicit adaptation through visual mechanisms, then Target Jump and Target Size manipulations may modulate implicit adaptation to different degrees. For instance, if task success perturbations affect implicit adaptation by changing the relative distance between the cursor and the center of the target, then Target Jump manipulations may be more effective than Target Size manipulations. Alternatively, the dynamic stimuli of Target Jump manipulations may affect implicit adaptation by drawing attentional resources (13), causing it to suppress adaptation more robustly than the less disruptive Target Size paradigm. Considering the increasing interest in the influence of TE and reward on motor learning and implicit adaptation [for review see (17, 18)], it is important to assess and understand the efficacy of these different task success manipulations.

In this report, we investigate how Target Size and Target Jump task success manipulations influence implicit adaptation. We first set out to replicate previous findings employing these perturbations. Our first two experiments investigate the effectiveness of Target Size manipulations. After some difficulties replicating this effect (Experiments 1-2), we employed Target Jumps to verify that task success exerts any influence on implicit adaptation (Experiment 3). Finally, we employed a wide range of errors in order to probe the efficacy and reliability of task success effects using both Target Size and Target Jump manipulations (Experiment 4). We find that Target Jump manipulations reliably and robustly influence implicit adaptation, more consistently than Target Size manipulations.

## MATERIALS AND METHODS

### Participants

Participants (*n* = 204, 124 female, 20.10 ± 1.72 years of age ranging from 18 to 29 years, 190 right-handed and 12 ambidextrous as determined by the Edinburgh Handedness Inventory [19]) were recruited from the Princeton University community. Participants who filled out demographic forms (*n* = 87) reported belonging to the following racial categories at the following rates: White – 48.3%, Asian – 32.2%, Black – 8%, More than one race – 7%, Prefer not to say – 2.3%. In addition, 9.2% of these participants reported being Hispanic or Latino, 89.6% reported not being Hispanic or Latino, and 1.1% preferred not to specify their ethnicity.

All participants provided informed, written consent in accordance with procedures approved by the Princeton University Institutional Review Board. Participants received either course credit or a $12 honorarium as compensation for their time. A power analysis (GPower V3.1) of Kim and colleagues’ (11) Experiment 1 indicated that 17 subjects per group would be required for 80% statistical power given their reported effect size (*Cohen’s d*=1.01), so we opted to collect 24 participants per group in Experiments 1 and 2, 50% larger than the initial study’s sample size (16/group). A power analysis of the results of Experiment 3 reported here indicated that 40 participants would be required to obtain sufficient statistical power to observe an effect of jumping the target provided the number of pre-planned post-hoc comparisons, and we collected data from 42 participants for Experiment 4 as a result of open study enrollment. Sample sizes for Experiment 3 (*n* = 18) were not determined by power analysis, but are greater than sample sizes in other studies in the literature investigating the effect of task success on implicit adaptation in the laboratory (11, 13).

### Apparatus

Participants performed a center-out reaching task while vision of the hand was obscured by an LCD monitor (60 Hz, 17-in., Planar Systems, Hillsboro, OR) mounted 27 cm above a digitizing tablet (125 Hz, Wacom Intuos Pro L, Wacom, Vancouver, WA). Participants controlled a visually-displayed cursor by moving a stylus, which was embedded in an air hockey paddle, with their right hand (**Fig. 1A**). We opted to use the air hockey paddle system as opposed to the stylus alone 1) to encourage participants to make arm movements about the shoulder and elbow joints instead of the joints of the wrist and fingers and 2) to replicate the experimental conditions of Kim et al. (11) as closely as possible (personal communication). Experimental software was programmed in Matlab R2013a using the Psychtoolbox extension V3.0, and was run on a Dell OptiPlex 7040 computer (Dell, Round Rock, TX) with a Windows 7 operating system (Microsoft Co., Redmond, WA). All stimuli were presented on a black background that filled the display. Experiments were conducted with the room lights extinguished to limit peripheral vision of the arm and to maximize stimulus visibility.

**Figure 1.**
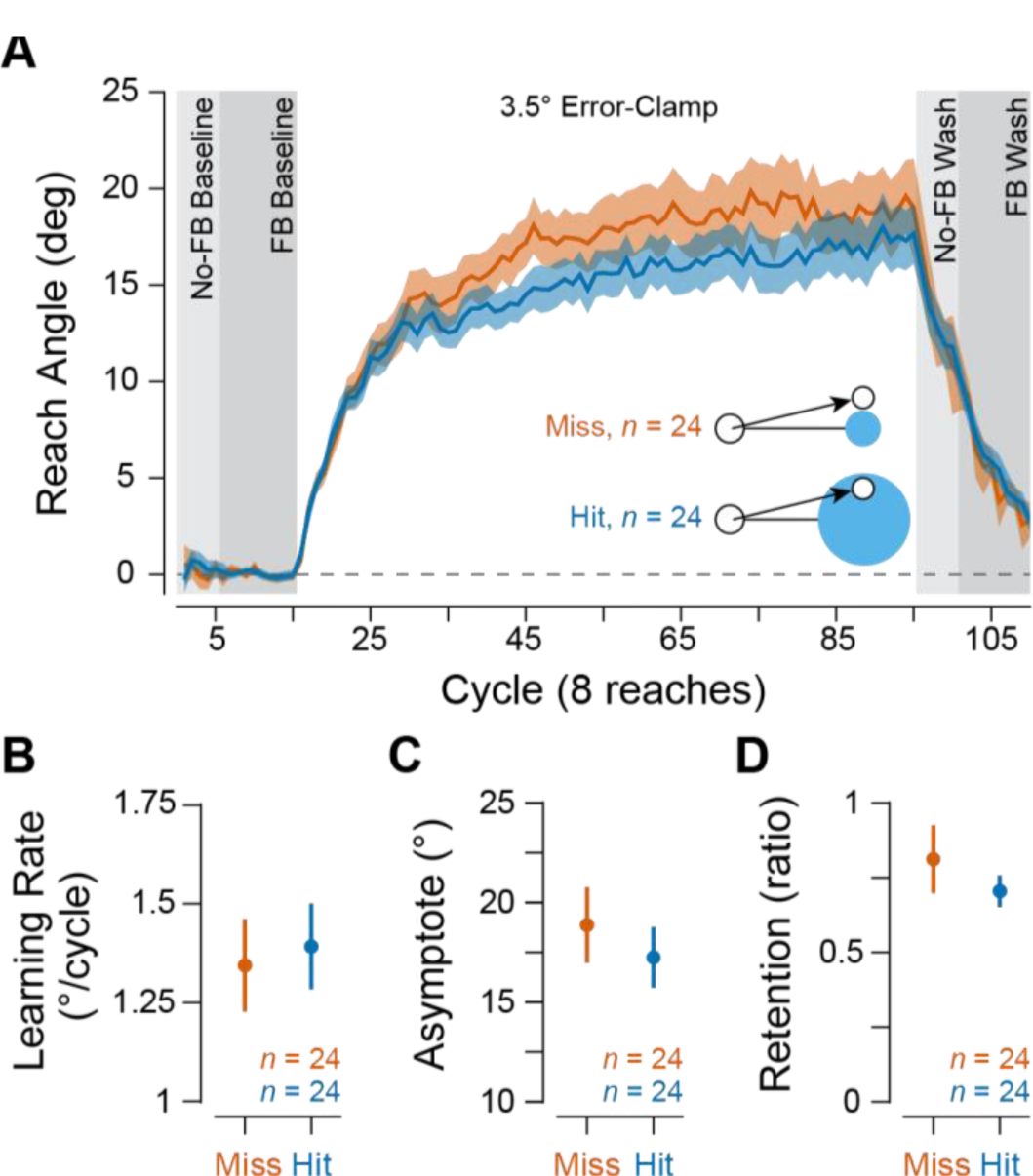
Effects of Target Size-based manipulations on implicit adaptation in Experiment 1. (**A**) Learning curves during Experiment 1. Participants in both the Miss (orange, *n* = 24 [16 female, 8 male]) and Hit (blue, *n* = 24 [16 female, 8 male]) groups exhibited robust changes in hand angle in response to the error-clamp perturbation. (**B**) Early learning rates during Experiment 1. Learning rate was quantified as the mean change in reach angle per cycle across the first 5 cycles of the experiment. (**C**) Asymptotic learning during Experiment 1. Asymptotic learning was quantified as the mean reach angle across the last 10 cycles of the error-clamp block. (**D**) Retention during the no-FB washout block in Experiment 1. Retention was quantified as the ratio of reach angle in the final cycle of the no-FB washout block to the reach angle in the final cycle of the error-clamp block. Data are shown as mean ± standard error of the mean. Abbreviations: FB, feedback.

### Cursor feedback

A visually-displayed cursor (filled white circle, 1.5 mm diameter in Experiment 4, 3.5 mm diameter in Experiments 1-3) provided movement-related feedback (FB), which was displayed with a 44 ms latency. Note that this latency was not deliberately programmed into the task, but a result of lag derived from tablet sampling rates, monitor refresh rates, and software/operating system processing time. During baseline and washout trials, the cursor either faithfully showed participants’ hand locations throughout the trial (FB trials) or was not displayed (no-FB trials). On “error-clamp” trials, the angle of the cursor was fixed off-target and participants could only control the radial distance of the cursor (**Fig. 1B**) (15). In combination with instructions to ignore the error-clamp FB and reach straight for the target, this manipulation reliably isolates implicit adaptation and minimizes explicit re-aiming (15, 20).

### Center-out reaching task

To initiate a trial, participants positioned the hand in a central start location (6 mm diameter) using a guide circle that limited cursor feedback between trials (radius = distance between the hand and the starting location). When the hand was within 1 cm of the start location, the guide circle disappeared and veridical cursor FB was displayed. After the hand was in the starting location for 500 ms, a blue (RGB blue) target appeared 8 cm away. Participants were instructed to quickly slice through the center of the target without stopping before returning to the start location to initiate the next trial. When provided, cursor FB at the target distance was sustained for 50 ms. If the target-directed movement duration exceeded 600 ms, “Too Slow” was displayed in red on the screen and played through the computer speakers after the trial. When the target was presented at multiple locations within an experiment, trials were presented in “cycles,” such that all targets were experienced at all possible locations before being repeated.

### Procedure

Task procedures for Experiments 1-4 are described in brief below. A document with the instructions delivered to participants can be found at the GitHub repository for this manuscript (https://github.com/kimoli/TaskSuccessImplicitAdapt).

#### Experiment 1

Experiment 1 aimed to faithfully replicate Experiment 1 of Kim et al. (11), which demonstrated attenuation of implicit motor learning when participants saw cursor FB that hit the target.

The session proceeded as follows: 5 cycles of no-FB baseline, 10 cycles of veridical-FB baseline, a 3-trial 45° clamp tutorial, 80 cycles of 3.5°-error-clamped FB (clamp direction counterbalanced across subjects), 5 cycles of no-FB washout, and finally 10 cycles of veridical-FB washout. During each cycle, targets appeared once in each of 8 possible locations: 0°, 45°, 90°, 135°, 180°, 225°, 270°, and 315°. The 3-trial clamp tutorial phase aimed to inform participants about the nature of the clamp through practice. On each trial, the target appeared straight ahead (90°), and participants were instructed to reach in different directions away from the target to demonstrate the lack of contingency between reach and cursor FB directions (Tutorial trial 1: straight to the right, trial 2: straight left, trial 3: straight back/towards the body). Following the tutorial, the experimenter instructed participants to ignore the cursor and try to slice through the center of the target with their hand.

Participants (*n* = 48) were divided into two groups. One group saw a larger, 16 mm diameter target, such that, during clamp trials, the cursor landed completely within the target (“Hit” group). The other group saw a smaller, 6 mm diameter target that excluded the error-clamped cursor (“Miss” group; **Fig. 1C**).

#### Experiment 2

Experiment 2 was designed to standardize participants’ perceptions of TE, regardless of visual FB, by employing tones to indicate success or failure. This experiment proceeded largely as described for Experiment 2, with the exceptions described below.

In addition to visual FB, participants (*n* = 96) received auditory FB at the end of each reach. A pleasant dinging sound played at the end of the trial when the cursor (or hand, during no-FB blocks) landed within a certain angular distance of the center of the target. Otherwise, an unpleasant knocking sound was played. Participants (*n* = 96, 24/group) were divided into 4 groups. The larger 16 mm diameter target was displayed to two groups (“Hit” groups) and the smaller 6 mm diameter target was displayed to the other two groups (“Miss” groups). Hit and Miss groups were further divided into groups with a stricter distance threshold for playing the pleasant dinging sound (6 mm, “Strict”) or a more lenient distance threshold (16 mm, “Lenient”), such that participants in the Strict groups heard the unpleasant sound at the end of each trial during the error-clamp block while participants in the Lenient groups heard the pleasant sound at the end of each error-clamp trial.

During the 3-trial, 45°-error clamp tutorial, in addition to instructions related to the cursor feedback, participants were instructed that the sounds would no longer correspond to their actual performance, and instead corresponded to the distance of the cursor relative to the center of the target. Thus, they had no control over both the trajectory of the clamped cursor and the sounds that would play at the end of the trial. Nonetheless, they were instructed to ignore the clamped feedback and still try to bring their hand through the center of the target. When participants reached the washout phases, they were informed that the auditory and cursor feedback once again reflected their performance.

#### Experiment 3

Experiment 3 was designed to test whether an alternative method of manipulating task success – the Target Jump – would effectively influence implicit adaptation. Since these effects have been reported previously, we modeled our study after experiments described by Tsay et al. (13).

Participants (*n* = 18) reached to a single target location (90° [straight ahead]) throughout the study. First, they performed 100 baseline trials during which they received veridical FB. Then, we explained the nature of the error-clamp manipulation to participants and walked them through 3 demonstration trials. These demonstration trials proceeded largely as described for Experiment 1 (above), except that 1) participants also saw the target jump to the final cursor location on the second demonstration trial, and 2) participants saw the target jump 45° past the final cursor location on the last demonstration trial. After the demonstration, we instructed participants to do their best to ignore both the error-clamped FB and the Target Jumps. Instead, we asked them to try to reach directly through the target’s initial location.

Subsequently, participants were exposed to 800 trials with 4° error-clamped FB, and the direction of the error-clamp was varied randomly on each trial to maintain mean levels of adaptation around zero during the study. Then, on each trial, participants saw one of four different possible target jump contingencies: No Jump, Jump-To, Jump-Away, and Jump-in-Place. As a control, on “No Jump” trials, the target simply appeared and underwent no changes during the trial. To test the effects of eliminating TE on implicit adaptation, “Jump-To” trials were included where the target jumped 4° so that the error-clamped cursor FB landed directly on the center of the target. To test the effects of increasing task error on implicit adaptation, on “Jump-Away” trials the target jumped 4° in the direction opposite the error-clamp so that the center of the target was 8° from the center of the error-clamped FB. Finally, “Jump-in-Place” trials on which the target was extinguished for 1 frame before being re-illuminated in the same location were included to control for attentional effects of the target disappearing from its original location. We opted to hide the target for a single frame, as this was the duration specified by an earlier report utilizing the jump-in-place manipulation (13). In our case, a single frame lasted 16.7 ms. This Jump-In-Place manipulation produced a noticeable change in the visual display that approximates the experience of noticing the displacement of the target in the Target Jump conditions. All target manipulations were implemented when the hand passed 1/6 of the distance to the target on each trial. Single-trial learning was quantified as the change in reach angle between two subsequent trials.

#### Experiment 4

Experiment 4 was designed to test whether the effects of task success on implicit adaptation depend on error magnitude and task success manipulation. Thus, we employed the Target Jump and Target Size manipulations, similar to what was described for Experiments 1-3, and measured single-trial learning as described for Experiment 3.

As in Experiment 3, all targets appeared straight ahead (90°), and the study began with a 100-trial baseline period with veridical cursor FB followed by a 3-trial error-clamp tutorial phase. Then, an 865-trial error-clamp phase began. During this phase, participants encountered error-clamp magnitudes of 1.75°, 3.5°, 5.25°, 7°, 8.75°, and 10.5° (clockwise and counterclockwise). On each trial, they also experienced one of three levels of task success: Miss, Hit, and Target Jump-To (Jump-To). On Hit trials, the target was 31 mm in diameter and completely encompassed the cursor on the 10.5° error-clamp trials. On Miss trials and Jump-To trials, the target was 4.5 mm in diameter and completely excluded the cursor on the 1.75° error-clamp trials. During Jump-To trials, the target shifted 1/6 of the way through the participant’s reach such that the cursor and target were concentric at the end of the trial.

#### Statistical Analysis

Raw data were preprocessed in MATLAB 2020a before being further processed and undergoing statistical analysis in R (RStudio, 1.3.959; RStudio, PBC, Boston, MA, R, 4.1.1). Because differences in approaches to data analysis may cause follow-up studies to fail to replicate initial reports, we analyzed the data following the approaches used in the studies we intended to replicate. Thus, for experiments solely dealing with Target Size task success manipulations (Experiments 1 & 2), we employed the approach described by Kim et al. (11) and measured reach angle at the hand position at the time of maximum velocity on each trial. For experiments including Target Jump manipulations (Experiments 3 & 4), we used the approach of Tsay and colleagues (13) and measured reach angle as the hand position at the time that the hand passed the center of the target. Two criteria were used to exclude trials from further analysis, based on the practices in the previous reports. First, trials on which the reach angle deviated from the target angle by more than 90° were excluded. Second, trials on which the reach angle deviated from the running average (5-trial window) by more than 3 standard deviations were also excluded. Across this report, <1% of trials were excluded (Experiment 1: 1%, Experiment 2: 0.6%, Experiment 3: 0.5%, Experiment 4: 1%) via these criteria. For Experiments 3 and 4, we also excluded trials on which participants reached toward the only/expected target location (straight ahead) before the target appeared. This led us to exclude an additional 3.7% of trials from Experiment 3 and 4.4% of trials from Experiment 4. For Experiments 1 & 2, veridical FB baseline biases for each participant at each target were then computed and subtracted from the reach angles. For Experiments 1 and 2, reach angles were subsequently binned by cycle (see *Procedure* above). Early learning rates were calculated as the estimated average change in hand angle over the first five cycles of the clamp block. To stably estimate the level of adaptation at cycle 5, cycles 3-7 were averaged. Asymptotic adaptation was estimated as the average reach angle over the last ten cycles of the clamp phase. Retention ratios were quantified as the ratio of reach angle in the final cycle of the no-FB washout phase to the reach angle in the final cycle of the preceding error-clamp phase. In the interest of replication (21), these definitions of learning rate, asymptotic performance, and retention were chosen for consistency with the report from Kim and colleagues (11).

In Experiments 3 and 4, single-trial learning was quantified as the difference in reach angle between subsequent trials. Individual participants’ performance within each trial type was averaged within clamp direction, and then these mean values were averaged. Finally, performance within trial type was compared across participants.

When comparisons were only made between two conditions for an experiment, we used Student’s t-tests (paired or unpaired, as was appropriate the sampling conditions). When comparisons were made between three or more conditions, we used a two-way ANOVA (repeated measures ANOVA was applied when appropriate for the sampling conditions). If main effects or interactions were found to be statistically significant in the ANOVA, we followed up with appropriate post-hoc comparisons. Type-I errors were limited by adjusting p-values to control the false-discovery rate.

### Code and Data Availability

Data and analysis code have been deposited at https://github.com/kimoli/TaskSuccessImplicitAdapt (https://doi.org/10.5281/zenodo.7982916).

### RESULTS

#### Experiment 1: Does manipulating task outcome via target size influence implicit motor learning?

Previous works have suggested that implicit adaptation can be modulated by task success cues, such as hitting a target (9, 11, 13). Before probing the conditions that drive this sensitivity to task success, we sought to replicate these prior findings. In Experiment 1, we replicated the approach and conditions of Kim and colleagues’ (11) first experiment, including employing the same 3.5° error-clamp size (see *Methods* for additional details). To maximize the likelihood that we would observe an effect, we tested the two most distinct task success conditions: Hit, (cursor lands completely inside the target, **Fig. 1A** inset, bottom), and Miss (cursor never touched the target, **Fig. 1A** inset, top). Based on a power analysis of the differences between asymptotic performance in Kim et al.’s (11) Miss and Hit groups, we included 24 participants in each group (total *n* = 48; see *Methods* for details of the power analysis).

Both the Hit and Miss groups showed substantial adaptation of reach angles opposite the direction of the error-clamp (**Fig. 1A**). However, we did not observe statistically significant effects of task success on early learning rates (Student’s two-sample t-test, *t*(46) = −0.30, *p* = 0.77, **Fig. 1B**) or asymptotic learning (*t*(46) = 0.67, *p* = 0.51, **Fig. 1C**), or retention (*t*(46) = 0.85, *p* = 0.40, **Fig. 1D**). Although the degree of adaptation exhibited by the Hit group (mean ± SD, 17.24° ± 7.46°) was numerically lower than that of the Miss group (18.88° ± 9.33°), the difference between group mean asymptotes observed here corresponds to a small effect size (*Cohen’s d* = 0.08) – much smaller than very large effect size seen by Kim and colleagues (11).

The aforementioned analysis used the analysis procedures of Kim and colleagues (11) and could not detect any significant effects of Target Size on adaptation. However, a trend can be seen where mean adaptation in the Miss group is numerically greater than adaptation in the Hit group for the entire error-clamp block. As an exploratory, post-hoc test, we compared performance averaged over the entire block but still found no statistically significant differences in the degree of adaptation (unpaired t-test, *t*(46) = 1.05, *p* = 0.30). In one final test, we compared performance during the error-clamp cycle exhibiting the greatest differences between the Miss and Hit groups (cycle 45), but still could not detect any significant differences (*t*(46) = 1.69, *p* = 0.10).

Therefore, our data suggest either that manipulating task success via target size does not affect implicit adaptation, or that the magnitude of the effect is much smaller than previously reported. This latter interpretation is consistent with the possibility of a “Decline Effect”, where an initially reported effect size is larger than those observed later (21). However, it is also possible that individual participants’ interpretations of what degree of cursor accuracy constitutes “good performance” may affect subjective experiences of task success during the error-clamp manipulations. The same error magnitude may be subjectively evaluated as “good enough” by one participant but as a “grave error” by another (22). In this case, differences in participants’ subjective evaluations of task success between our sample and the sample collected by Kim et al. (11) may account for differences in our results. To address this, we conducted another experiment that included auditory cues to clarify the task success conditions to participants and explicitly demarcate task success from task failure using auditory rewards.

#### Experiment 2: Does clarifying task success conditions with auditory feedback reveal an effect of task success on implicit adaptation?

To address the possibility that Target Size differences alone failed to affect participants’ perceptions of task success during Experiment 1, we provided additional, auditory task success cues in Experiment 2. As in Experiment 1, participants (*n* = 96) either reached to a large target that encompassed the clamped cursor feedback (Hit) or a small target that excluded the clamped cursor feedback (Miss). In addition to these visual task success FB cues, we played auditory FB at the end of the movement.

During the baseline and washout periods, auditory cues were contingent upon hand position at the end of the trial, thereby establishing an association between the auditory FB and participants’ perceptions of task success. For participants assigned to “Strict” auditory FB conditions, a pleasant chime sound was played if the hand landed within the radius of the smaller possible target size, regardless of the displayed target size. Otherwise, an unpleasant knocking sound was played. In contrast, participants assigned to the “Lenient” auditory FB conditions heard the pleasant chime sound when the hand landed within the radius of the larger possible target size. At the onset of the Error-Clamp block, auditory FB became contingent upon the error-clamped cursor FB instead of the hand position such that participants assigned to “Lenient” conditions heard the pleasant chime sound during the 3.5° error-clamp phase whereas participants assigned to the “Strict” conditions heard the unpleasant knocking sound (**Fig. 2A**).

**Figure 2.**
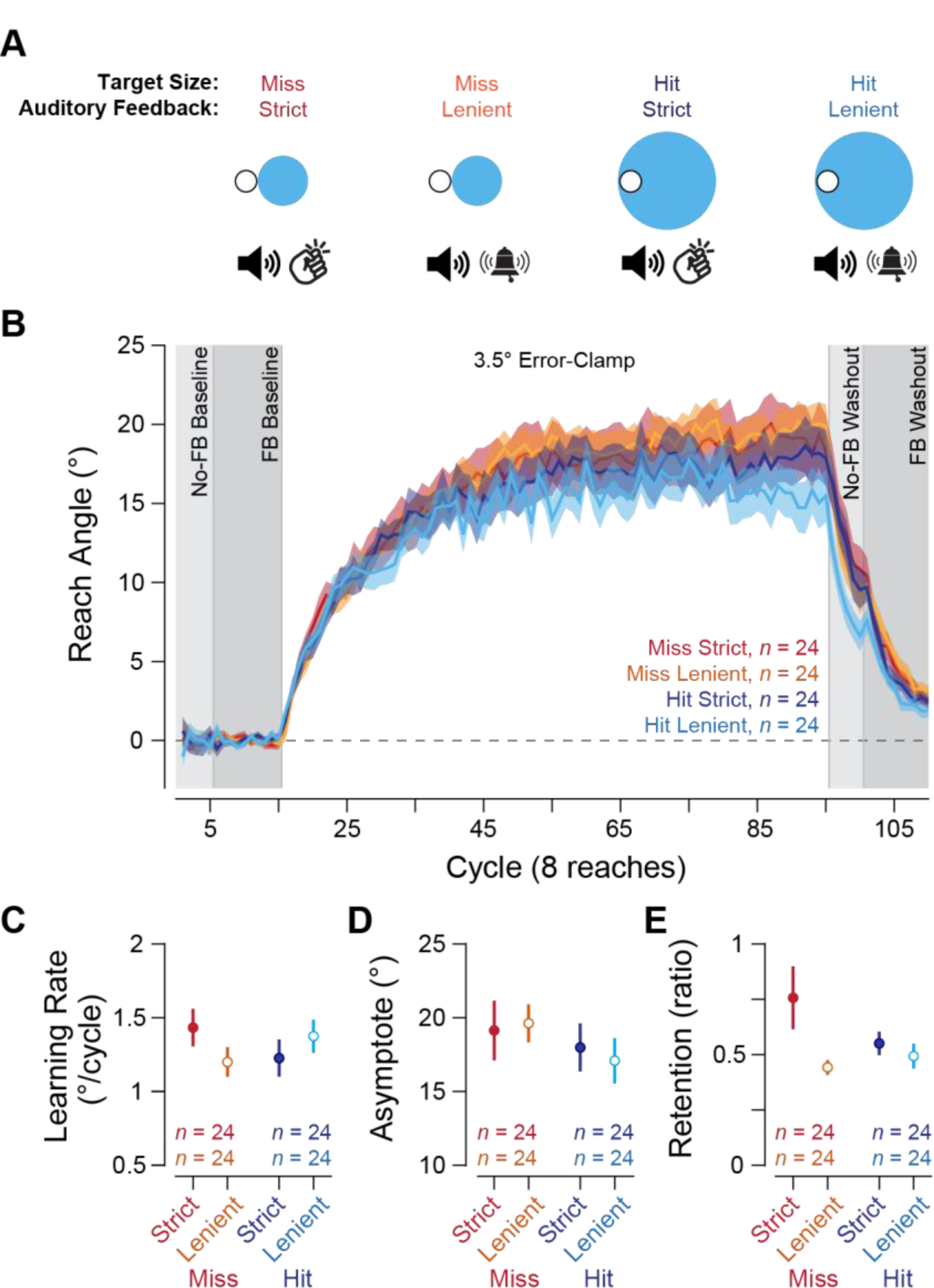
Effects of manipulations of task success using auditory cues in Experiment 2. (**A**) Schematic of visual FB and auditory cues presented to participants during the Error-Clamp block. In Strict conditions (first and third configurations), a knock sound played when the 3.5° error-clamped FB reached the target distance, regardless of target size. In Lenient conditions (second and third configurations), a pleasant dinging sound was played instead. (**B**) Learning curves during Experiment 2. All groups exhibited robust learning in response to the error clamp (*n* = 96 [45 female, 44 male, 7 prefer not to say]). (**C**) Early learning rates during Experiment 2. (**D**) Asymptotic learning during Experiment 2. (**E**) Retention ratios during washout of Experiment 2.

Participants were divided into 4 equally-sized groups according to a 2 x 2 design with 2 levels of auditory cue condition (Strict or Lenient) and 2 levels of task success condition (Hit or Miss, as in Experiment 1). This design allowed us to systematically test whether adding auditory reward and punishment FB to visual indicators of task success would reveal an effect of task performance on implicit adaptation. If auditory FB effectively enhances participants’ experiences of task success and task success suppresses implicit adaptation, then participants in the Hit Lenient condition ought to have shown significantly lower levels of asymptotic adaptation relative to participants in the Miss Strict condition.

First, to confirm that participants correctly interpreted the pleasant and unpleasant auditory cues as indicating task success, we examined how participants adjusted their reach angle in response to the auditory FB in the No-FB baseline phase (when they were encouraged to hit the target). During trials with reach endpoints in the range where the tone played varied between groups (*i.e.*, between the small and large target diameters; **Fig. 2A**, left), there was a significant effect of auditory FB condition (two-way between-subjects ANOVA, *F*(1,92) = 4.16, *p* = 0.04, *partial η^2^* = 0.04) but not target size (*F*(1,92) = 0.72, *p* = 0.40) or the interaction between the two factors (*F*(1,92) = 0.02, *p* = 0.88) on changes in hand angle on the subsequent trial. A post-hoc t-test confirmed that adjustments in reach angle were greater among participants in the Strict groups (mean ± standard deviation: 4.82 ± 1.53°) compared to those in the Lenient groups (4.21 ± 1.40; *t*(94) = 2.05, *p* = 0.04, *Cohen’s d* = 0.42), indicating that the auditory cues were understood by the participants to indicate success or failure.

Notably, when cursor FB was provided alongside veridical cursor FB in a subsequent baseline phase, auditory FB ceased to influence the magnitude of updates to reach angle within the analyzed window (*F*(1,92) = 0.93, *p* = 0.34), and target size drove differences between groups (*F*(1,92) = 7.74, *p* = 0.007, *partial η^2^* = 0.08) without an interaction between the factors (*F*(1,92) = 0.43, *p* = 0.51). A post-hoc t-test showed that updates were significantly larger among participants in Miss conditions (small target; mean ± SD: 3.70 ± 0.83°) than those in Hit conditions (large target; 3.19 ± 0.96°; *t*(94) = 2.79, *p* = 0.006, *Cohen’s d* = 0.57). These findings suggest that, when available, visual indicators of task success take precedence in guiding explicit performance over other modalities of performance FB.

During the error-clamp phase, when auditory FB was clamped alongside cursor FB and participants were instructed not to re-aim their movements based on the FB they received, all groups exhibited robust learning to the error clamp (**Fig. 2C**). However, auditory cues, target size, and their interaction had no effect on participants’ learning rates (**Fig. 2D**) or asymptotic levels of adaptation (**Fig. 2E;** refer to **Table 1** for details of statistical tests). Thus, even with the addition of auditory cues, task success indicators did not effectively modulate the acquisition of implicit motor adaptation.

**Table 1.** Details of two-way between subjects ANOVAs conducted for Experiment 2.

| Factor | F | df<sub>n</sub> | df<sub>d</sub> | p |
| --- | --- | --- | --- | --- |
| Early Learning Rate |  |  |  |  |
| Strict/Lenient Auditory FB | 0.13 | 1 | 92 | 0.72 |
| Hit/Miss Target Size Condition | 0.02 | 1 | 92 | 0.89 |
| Auditory x Target Size Interaction | 2.66 | 1 | 92 | 0.11 |
| Asymptotic Adaptation |  |  |  |  |
| Strict/Lenient Auditory FB | 0.02 | 1 | 92 | 0.90 |
| Hit/Miss Target Size Condition | 1.27 | 1 | 92 | 0.26 |
| Auditory x Target Size Interaction | 0.18 | 1 | 92 | 0.67 |

*Note.* Abbreviations: df, degrees of freedom.

During the No-FB washout phase, auditory FB significantly affected retention of implicit adaptation (two-way between subjects ANOVA, *F*(1,92) = 5.06, *p* = 0.03, *partial η^2^* = 0.05) while Target Size (*F*(1,92) = 0.88, *p* = 0.35) and the interaction (*F*(1,92) = 2.41, *p* = 0.12) had no effect on retention (**Fig. 2F**). A post-hoc t-test indicated that retention was greater among participants in the Strict condition (mean ± standard error: 0.65 ± 0.08 retention ratio) than participants in the Lenient condition (0.47 ± 0.03 retention ratio; two-sample t-test, *t*(94) = 2.23, *p* = 0.03, *Cohen’s d* = 0.46). This suggests that auditory FB may influence the rate of decay of implicit adaptation. However, we note that participants in the Lenient auditory conditions experienced an abrupt shift from hearing the pleasant to the unpleasant tone at the onset of the washout block when auditory feedback was released from the clamp perturbation and became contingent on reach angle. Indeed, many participants in the Lenient group noted the abrupt change and verbally questioned the experimenter about it, but this was not the case for the Strict group. So, it is unclear whether retention of implicit adaptation was suppressed by exposure to the pleasant tone during training, or whether performance in the Lenient conditions was disrupted by an auditory startle response or re-aiming in an attempt to control the auditory FB.

Notwithstanding a potential effect of auditory reward FB on retention in the washout phase (potentially mediated by explicit re-aiming), the addition of performance-related auditory cues did not substantially affect the rate or degree of implicit adaptation. This is in line with the results of Experiment 1, providing further evidence that manipulating task success using different target sizes does not affect implicit adaptation, or the effect is quite small. Additionally, auditory cues of task success may not significantly impact implicit adaptation, consistent with previous work showing that auditory cues may primarily shape explicit motor learning (23). Taken together, the results of these first two experiments suggest that the effect of manipulating task success via changes in target size is either small or nonexistent. Thus, we sought to assess whether another method for manipulating task success – the so-called “Target Jump” after the fashion of Leow and colleagues (9) and Tsay and colleagues (13)–influences implicit adaptation.

#### Experiment 3: Do task success manipulation using Target Jumps influence implicit motor learning?

In Experiment 3, we aimed to replicate recent work employing a different form of task success manipulation – the Target Jump – that demonstrated an effect on single-trial learning (STL) (13). During Target Jump manipulations, the target is displaced partway through the trial so that the cursor feedback lands at an experimenter-specified distance from the center of the target (**Fig. 3A**, top), thereby manipulating task success without manipulating the size of the target. As Target Jumps have been shown to modulate learning in block designs (9, 10), we suspected that replications of an effect of jumping the target may prove more forthcoming.

**Figure 3.**
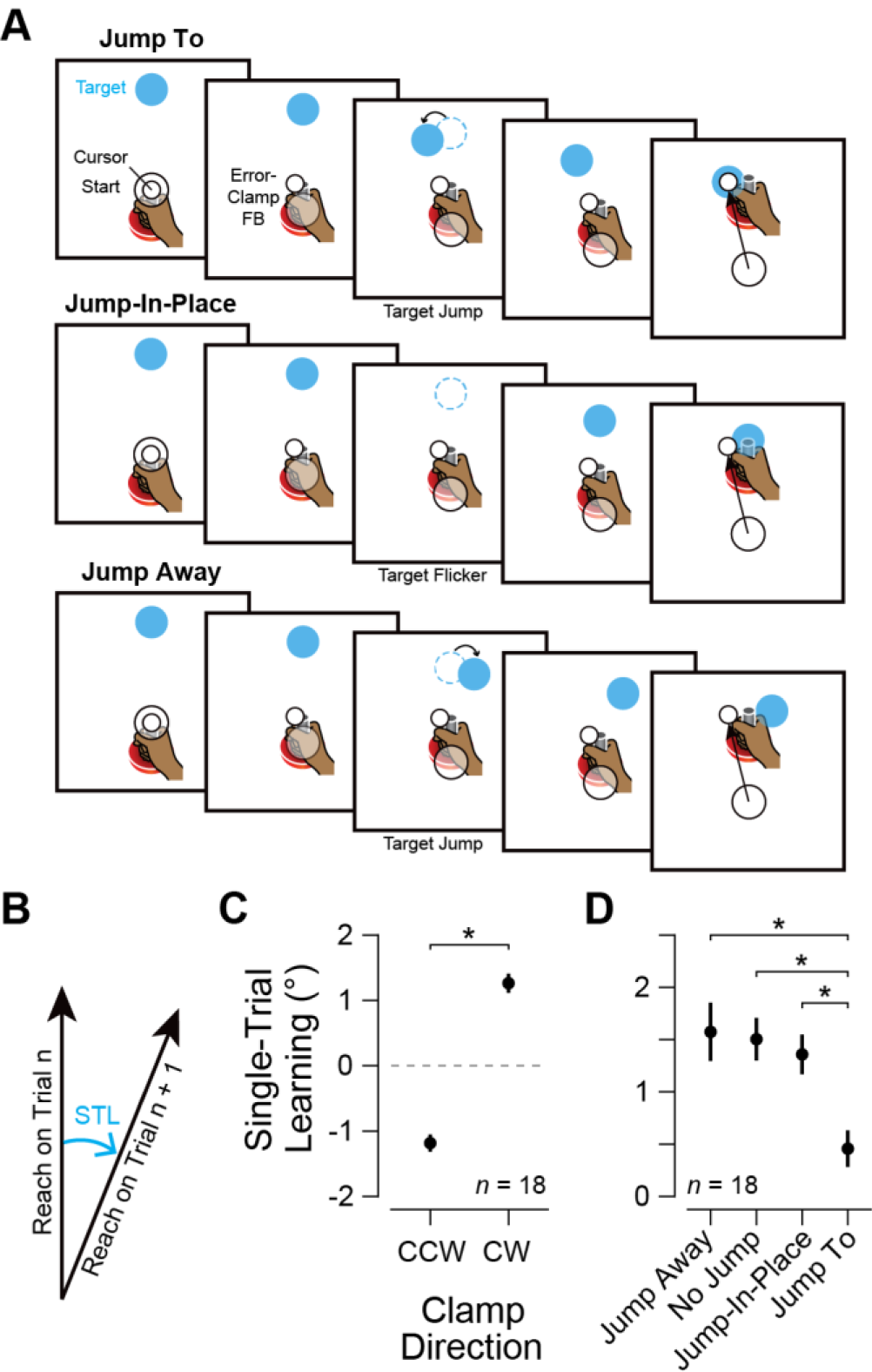
Effects of Target Jump manipulations on single-trial, implicit adaptation. (**A**) Schematic illustrating the different Target Jump perturbations. (**B**) Schematic showing how single-trial learning (STL) was computed for this experiment. (**C**) STL in response to either counterclockwise or clockwise error-clamped FB. Positive STL indicates a counterclockwise change in reach angle, while negative STL indicates a clockwise change. * indicates a statistically significant one-way within-subjects ANOVA outcome with *p* < 0.001 (*n* = 18 [8 female, 10 male]). (**D**) STL in response to 4° error-clamped cursor FB paired with the Target Jump manipulations indicated on the x-axis. For this panel, STL has been computed such that positive STL indicates adaptation in the direction opposite the error-clamp (*i.e.*, error-appropriate adaptation), and negative STL indicates adaptation in the same direction as the error-clamp. *s indicate paired-t-test adjusted p-values of 0.02. Abbreviations: FB – feedback, STL – single-trial learning.

Participants (*n* = 18) were instructed to reach directly for the target that appeared and ignore any deflections in cursor FB or movement of the target, after the fashion of Tsay et al. (13). After a baseline period with veridical FB, all trials provided 4° error-clamped FB and one of four possible target perturbation events halfway through each reach. The direction of the error-clamped FB (clockwise or counterclockwise) was randomly varied across trials to maintain an average background level of 0° of accumulated adaptation, and adaptation in response to each error/Target Jump combination on trial *n* was quantified as the difference in reach angles on trials *n* and *n + 1* (STL, **Fig. 3B**). “Jump-To” trials, where the target was displaced by 4° such that endpoint cursor FB would fall on the center of the target (**Fig. 3A**, top), were included to assess whether eliminating task error via a Target Jump would affect implicit adaptation. “Jump-Away” trials, where the target was displaced by 4° away from the direction of the error-clamp, were included to assess whether increasing task error via a Target Jump would affect implicit adaptation (**Fig. 3A**, middle). “Jump-In-Place” trials, where the target disappeared for one frame, were included to control for potential attentional effects of the disappearance of the target in Jump-To and Jump-Away trials (**Fig. 3A**, middle), Finally, “No-Jump” trials, where the target was not perturbed during the trial, were included to provide a baseline rate of learning.

Participants showed robust, direction-specific STL in response to error-clamped feedback (one-way within-subjects ANOVA, *F*(1,17) = 94.7, *p* <0.001, *partial η^2^* = 0.84; **Fig. 3C**) that was, as reported by Tsay et al. (13), affected by target manipulations (*F*(1.16, 19.8) = 8.80, *p* = 0.006, *partial η^2^* = 0.23). In line with the previous report, Jump-To target perturbations significantly suppressed adaptation relative to No Jump (paired t-test, *t*(17) = 3.36, *p_adj_* = 0.02, *Cohen’s d* = 1.43), Jump-Away (*t*(17) = 2.89, *p_adj_* = 0.02, *Cohen’s d* = 1.26), and Jump-In-Place trials (*t*(17) = 3.12, *p_adj_* = 0.02, *Cohen’s d* = 1.28, **Fig. 4D**). Contrary to the report by Tsay et al. (13), we did not observe a significant effect of Jump-In-Place perturbations on adaptation (*t*(17) = 1.96, *p_adj_* = 0.1). Given the lack of other significant differences between the conditions, the observation that only Jump-To target perturbations influence STL without attentional effects of Jump-In-Place trials or STL-enhancing effects of Jump-Away perturbations is not clearly consistent with graded effects of task success on implicit adaptation due to attentional distraction induced by the Target Jump.

**Figure 4.**
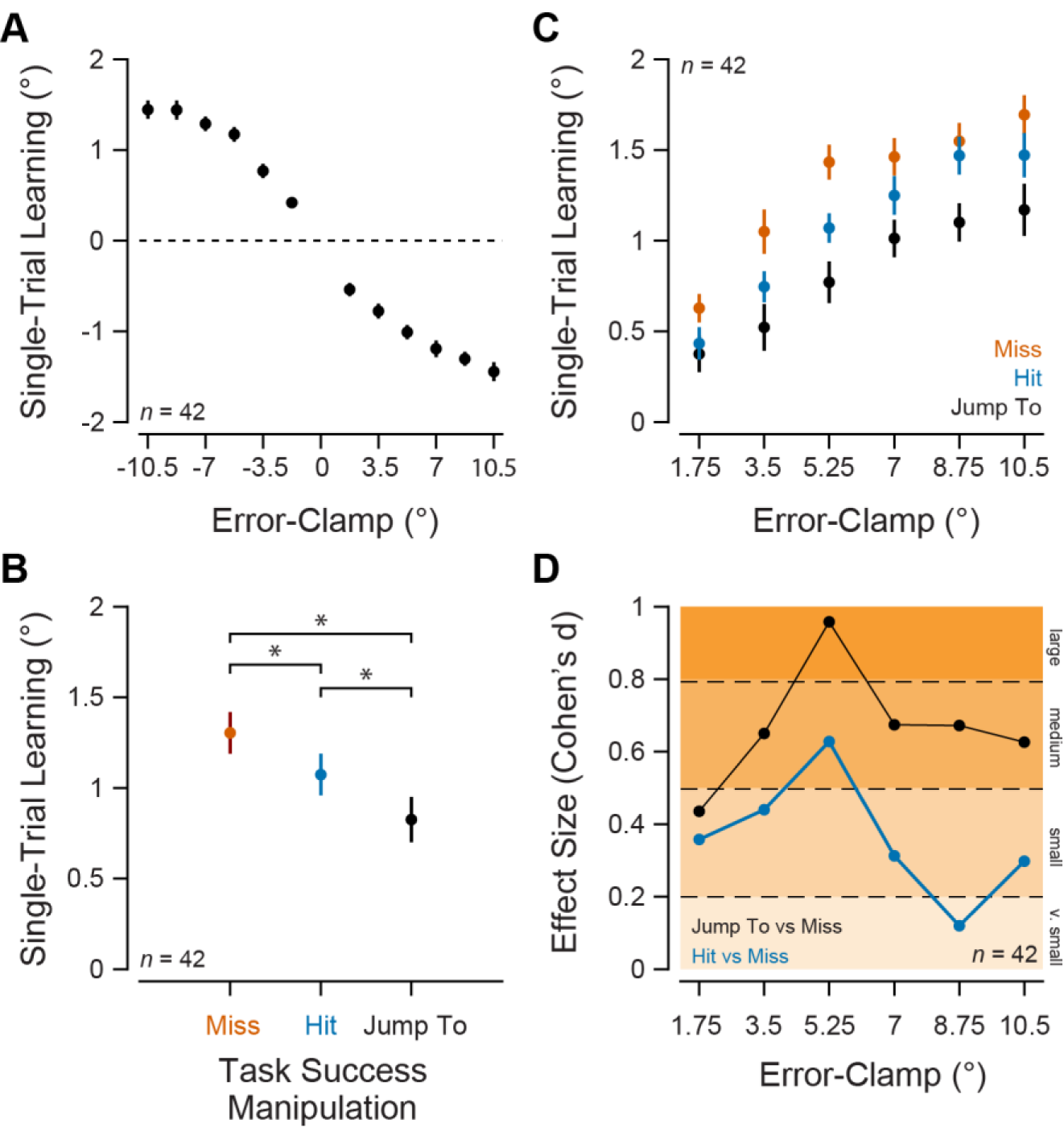
Effects of Target Size, Target Jump, and error-clamp size manipulations on single-trial, implicit adaptation. (**A**) STL in response to either counterclockwise or clockwise error-clamped FB, collapsed across task success conditions. Positive STL indicates a counterclockwise change in reach angle, while negative STL indicates a clockwise change (*n* = 42 [36 female, 5 male, 1 prefer not to say]). (**B**) STL collapsed across error-clamp magnitude/direction but separated by task success condition. Positive STL indicates a change in reach angle opposite the direction of the observed error-clamp. Boxplot center: median, box edges: 1st and 3rd quartiles, notch: 95% confidence interval of the median, whiskers: most extreme values not considered outliers. From left to right, *s indicate adjusted p-values from paired t-tests of <0.001, <0.001, and 0.02. (**C**) STL in response to error-clamped FB collapsed across direction but separated by error-clamp magnitude and task success condition. See **Table 2** for complete details on statistical comparisons. (**D**) Effect size measures (*Cohen’s d*) of the differences between the Miss condition and the Jump To (black) or the Hit (blue) conditions as a function of the magnitude of the error-clamp. Orange shading and labels on the right hand side of the panel indicate descriptions of effect sizes according to Cohen’s (24) threshold guidelines. Abbreviations: STL - single-trial learning.

The robust effects of Target Jumps on implicit adaptation replicated in Experiment 3 stand in stark contrast to the small-to-nonexistent effects of Target Size manipulations reported earlier in this manuscript. We noted, however, that the effect of target size was most pronounced in Experiment 3 of Kim and colleagues’ (11) report, which employed a smaller (1.75°) clamp. Thus, it is possible that the target size manipulation is strongest under conditions where errors are small. Notably, Target Jump manipulations may enjoy a degree of immunity to changes in error-clamp size as their effects have been observed with perturbations upwards of 30 degrees (9, 13). Experiment 4 addresses these hypotheses.

#### Experiment 4. How do Target Size and Target Jump manipulations influence implicit motor learning at various error sizes?

Experiment 4 employed multiple error-clamp sizes, Target Size manipulations (as in Experiments 1 & 2), and the Target Jump manipulation (as in Experiment 3). This was done in order to comprehensively assay the effect of each manipulation at various error sizes, as we speculated that target size manipulations may only be effective at small clamp angles. Participants (*n* = 42) were instructed to move straight toward the target that appeared on the screen, regardless of cursor FB, which would be clamped away from the center of the target by an angular error that randomly varied on each trial between 1.75° and 10.5°, at increments of 1.75°. Additionally, on a given trial, the target would be a) small enough that even the 1.75° clamp would miss the target (Miss; **Fig. 1A** inset top), b) large enough that even the 10.5° clamp would be entirely within the target (Hit; **Fig. 1A bottom**), or c) the target, at the same size as the Miss target, would jump to meet the cursor FB, eliminating task error (Jump-To; **Fig. 3A top)**. Clamp direction (clockwise or counterclockwise) varied across trials with zero-mean, allowing us to measure single-trial learning (STL) as the change in hand angle on trial t+1 in response to the error observed on trial t.

Participants exhibited robust STL which tracked the error magnitude and direction (stats, **Fig. 4A**). A two-way repeated-measures ANOVA highlighted a statistically significant effect of error-clamp magnitude (*F*(3.66, 150.05) = 62.14, *p* <0.001, *η ^2^* = 0.20) and task success condition (*F*(1.61, 65.99) = 17.96, *p* <0.001, *η ^2^* = 0.08), but no interaction (*F*(10, 410) = 0.97, *p* = 0.47). STL was significantly suppressed relative to the Miss condition by both Hit (*t*(41) = 4.04, *p_adj_* <0.001^4^, *Cohen’s d* = 0.50) and Jump-To FB (*t*(41) = 5.36, *p_adj_* <0.001, *Cohen’s d* = 0.91), although STL was suppressed more by Jump-To FB than Hit FB (*t*(41) = 2.79, *p_adj_* = 0.02, *Cohen’s d* = 0.49; **Fig. 4B**).

Subsequent pre-planned post-hoc pairwise comparisons provided further evidence that the Jump-To manipulation generally suppressed STL more than the Hit manipulation. While participants exhibited significantly less STL on Jump-To trials than Miss trials for all error-clamp magnitudes greater than 1.75°, participants only exhibited less STL on Hit trials relative to Miss trials at 3.5° and 5.25° error-clamps (**Fig. 4C**, see **Table 2** for statistical details). In addition, STL was significantly lower in the Jump-To than the Hit condition at 5.25° and 8.75° error-clamps (**Fig. 4C**, **Table 2**). Moreover, differences in STL between Miss and Jump-To conditions exhibited larger effect sizes than differences between Miss and Hit conditions at all error-clamp magnitudes greater than 1.75° (**Fig. 4D**, **Table 2**).

**Table 2.** Details of pre-planned post-hoc pairwise comparisons conducted for Experiment 4 (comparisons in *Fig. 4C*)

| Task Success x<br>Error-Clamp A | Task Success x<br>Error-Clamp B | t | p | p<sub>adj</sub> | Signif. | Cohen's<br>d |
| --- | --- | --- | --- | --- | --- | --- |
| Miss x 1.75° | Hit x 1.75° | 1.62 | 0.11 | 0.13 |  | 0.36 |
| Miss x 3.5° | Hit x 3.5° | 2.39 | <0.001 | 0.04 | * | 0.44 |
| Miss x 5.25° | Hit x 5.25° | 3.49 | 0.001 | 0.003 | * | 0.63 |
| Miss x 7° | Hit x 7° | 1.80 | 0.08 | 0.12 |  | 0.31 |
| Miss x 8.75° | Hit x 8.75° | 0.74 | 0.5 | 0.49 |  | 0.12 |
| Miss x 10.5° | Hit x 10.5° | 1.84 | 0.07 | 0.12 |  | 0.30 |
| Miss x 1.75° | Jump To x 1.75° | 1.72 | 0.09 | 0.13 |  | 0.43 |
| Miss x 3.5° | Jump To x 3.5° | 3.12 | 0.003 | 0.008 | * | 0.65 |
| Miss x 5.25° | Jump To x 5.25° | 5.49 | <0.001 | <0.001 | * | 0.96 |
| Miss x 7° | Jump To x 7° | 4.07 | <0.001 | 0.001 | * | 0.67 |
| Miss x 8.75° | Jump To x 8.75° | 3.57 | <0.001 | 0.003 | * | 0.67 |
| Miss x 10.5° | Jump To x 10.5° | 3.93 | <0.001 | 0.001 | * | 0.63 |
| Hit x 1.75° | Jump To x 1.75° | 0.50 | 0.62 | 0.62 |  | 0.09 |
| Hit x 3.5° | Jump To x 3.5° | 1.38 | 0.17 | 0.19 |  | 0.32 |
| Hit x 5.25° | Jump To x 5.25° | 2.30 | 0.03 | 0.046 | * | 0.46 |
| Hit x 7° | Jump To x 7° | 1.66 | 0.1 | 0.13 |  | 0.35 |
| Hit x 8.75° | Jump To x 8.75° | 3.11 | 0.003 | 0.008 | * | 0.54 |
| Hit x 10.5° | Jump To x 10.5° | 1.67 | 0.1 | 0.12 |  | 0.35 |

*Note.* All comparisons' degrees of freedom are equal to 41. Familywise error rates were maintained by adjusting p-values according to the false discovery rate method, accounting for the comparisons made in **Fig. 4B-C**. Comparisons reaching statistical significance ( $p_{adj} < 0.05$ ) are flagged with a \*.

Taken together, the results of Experiment 4 illustrate that the Jump-To manipulation generally elicits a larger and more reliable suppression of STL than the Hit manipulation for nearly all error sizes. However, the data do not provide clear support for the claim that the effect of the Hit manipulation becomes weaker as the magnitude of the error-clamp increases. Overall, there is a slight reduction of Hit effect size with increases in error-clamp magnitude, but the jump in effect size at 5.25° degrees disrupts this trend (**Fig. 4D**). We note that there were no extreme outliers (STL beyond 3 standard deviations from the mean) in the Miss or Hit conditions driving this change in effect size.

### DISCUSSION

In sum, the findings reported here highlight methodological variations that influence the effect of task success on implicit adaptation, with two main conclusions. First, the data indicate that implicit adaptation of reach angle is robustly suppressed when the target is jumped to the final cursor location (Target Jump manipulations [9, 13]; **Figs. 3-4**), in line with previous work. Second, implicit adaptation is at most mildly suppressed when the target is large enough to encompass the final position of an error-clamped cursor (Target Size manipulations (11); **Figs. 1-2, 4**). This stands in contrast to a prior report with large effects of Target Size manipulations (11). Critically, our Experiments 1 and 2 employed sample sizes twice as large as those used by Kim and colleagues (11), suggesting that the earlier study may have been underpowered and overestimated the magnitude of Target Size effects on implicit adaptation (21). Supporting the conclusion that effects of Target Size are relatively small, Experiment 4 comprehensively explored the effects of Target Jump and Target Size manipulations over a range of SPE magnitudes. Together, our results indicate that Target Jumps are the more efficacious of the two task success manipulations, and that Target Size manipulations produce relatively small effects.

#### What does the difference in efficacy between Target Jump and Target Size manipulations tell us about the mechanisms underlying the effect of task success on implicit adaptation?

The difference between the efficacy of Target Size and Target Jump manipulations suggests that the suppression of learning in these task success paradigms is not fully attributable to vision of the cursor hitting the target. Below, we explore three possible sources of the larger effect sizes observed with Target Jump manipulations: 1) enhanced salience, 2) graded interpretations of task success, and 3) distracted attention.

One might argue the larger effect of Target Jumps is observed because Experiments 3-4 employed a pseudorandom trial order, enhancing the salience of the task success manipulation. In contrast, the block designs of Experiments 1-2 allowed the task success signal to be fully predicted, reducing the salience of the Target Size manipulation. This however would not explain the differences in effect sizes between Target Size and Target Jump manipulations observed in Experiment 4, but it does bring up an interesting point about the advantages of single-trial-learning paradigms. In addition to the salience of the manipulation being enhanced by the lack of predictability, single-trial-learning paradigms benefit from greater statistical power due to being a within-subjects design.

It seems reasonable to speculate that Target Size more weakly modulates adaptation because the cursor does not lie as close to the center of the target and cannot be interpreted as wholly successful. In line with this possibility, Tsay and colleagues have suggested that a change in target position during a Target Jump changes the participant’s experience of “task error” to modulate adaptation in a graded fashion (13). Our data do not fully support this explanation, but they also do not clearly refute it. We did not observe an increase in the amount of adaptation observed during Jump-Away trials, suggesting that increases in task error measured as the distance between the cursor and the center of the target do not exert a graded effect on adaptation (**Fig. 3**). However, considering that the amount of single-trial implicit adaptation observed saturated around error-clamp magnitudes of 5.25° in the Miss condition (**Fig. 4**), it is possible that we failed to observe an enhancement of adaptation in the Jump-Away condition with the 4° error-clamp used in Experiment 3 due to a ceiling effect. Thus, while our data are not wholly consistent with the idea that there is a continuous task error variable that modulates implicit adaptation in a graded fashion, it seems plausible that the Target Jump manipulation may be read as greater task success and thereby have a greater impact on implicit adaptation than Target Size manipulations.

Target Jumps may alternately influence adaptation more robustly than Target Size manipulations because they involve a salient change in the visual scene that draws attentional resources and detracts from adaptation to SPEs (13). However, we did not observe a detrimental effect of briefly removing the target (Jump-In-Place, **Fig. 4**) that would allow us to infer that enhanced adaptation was masked by jump-related-distraction in the Jump-Away condition. We note that our Jump-In-Place manipulation likely involved removing the target from view for a briefer period of time than that used by Tsay and colleagues (13), as our in-laboratory setup allowed us to limit the duration of 1 frame of blank time to 16 ms, whereas the online setup employed by Tsay and colleagues (13) may have resulted in greater blank time. So, we do not claim that Jump-In-Place manipulations do not influence implicit adaptation, simply that the duration of changes in the visual scene in our experiments were unlikely to drive attentional capture effects that can explain the differences in Target Jump and Target Size manipulation efficacy. Instead, differences in the extent of adaptation may be explained by diminishing error relevance as the new target location grows increasingly distant from the initial (planned) target location (25).

#### Mechanisms underlying effects of task success on implicit adaptation

How might task success information influence the neural substrate for implicit adaptation? Considering that visuomotor reach adaptation is impaired among patients with spinocerebellar ataxia [SCA; (15, 26)], it is tempting to speculate that task success ultimately affects implicit adaptation by influencing activity in the cerebellar circuit. However, many SCA subtypes are characterized by substantial extra-cerebellar degeneration in the basal ganglia and brainstem alongside the cerebellar degeneration associated with the disease (27). So, it is not a foregone conclusion that task success modulates implicit adaptation via activity in the cerebellum.

For a brain region to mediate the effects reported here, that region ought to process signals related to both task success and implicit adaptation. Although, to our knowledge, there are no examples in the literature of neuroimaging or neural recording studies probing both task success and isolating implicit reach adaptation, we can take some clues from work involving task success and reach adaptation in general. Because the implicit process proceeds unless deliberate measures are not taken to suppress it (15, 28–31), neural activity during general visuomotor reach adaptation tasks ought to include activity that supports the implicit process. From the neuroscience literature, we know that the cerebellum, basal ganglia, motor cortex, and parietal cortices are involved in reach adaptation tasks with and without target jumps (32-37). For example, Diedrichsen and colleagues (38) conducted an fMRI study presenting both target jumps and visuomotor rotations during a reaching task, and they reported that the cerebellum, motor cortex, and basal ganglia all exhibited changes in BOLD responses to both kinds of visual feedback. Interestingly, visuomotor rotations and target jumps causing task failure induced similar changes in activity in the cerebellum and motor cortex. This observation suggests that the cerebellum and motor cortex may encode TEs and SPEs in a similar fashion. However, we note that target jump-driven activity in Diedrichsen et al. (38) may also have been related to changes in reach goals that were encouraged during their task. Taking this into consideration, we cautiously speculate that conflicting signals from task success (reduced TE) and SPEs in the cerebellum and motor cortex may underlie the suppressive effect of task success on implicit adaptation. As the mechanism by which task success influences adaptation has yet to be specified, this topic remains an important frontier for future investigation.

Prior work has speculated that effects of task success on implicit adaptation are mediated by reward-related processes and systems (9, 11). While we cannot rule out this interpretation, we do not favor it, as our experiments with auditory reinforcements did not clearly affect implicit adaptation (**Fig. 2**). Although auditory cues affected our retention metric, these effects likely arose from re-aiming in response to the abrupt change in auditory cue (23) (see *Results: Experiment 2* for more details). Given the lack of other statistically significant effects on uncontaminated parameters, it appears that the auditory cues in Experiment 2 did not modulate implicit adaptation. Although motor learning in general is sensitive to auditory reward cues (23, 39, 40), our interpretation is consistent with prior studies that have not reported auditory cue-driven recalibration of the internal model (23, 39, 41). However, while simple tones may not directly influence implicit adaptation, reward cues with greater motivational salience may elicit an effect. Indeed, prior work has suggested that monetary rewards can enhance the effects of task success during motor skill training (42), although it is not clear that these effects are mediated by the implicit process isolated in the error-clamp paradigm used here. Motor learning, broadly, is sensitive to reward feedback (43-47), and future work providing rewards while isolating implicit adaptation will be important for fully addressing this issue.

#### Summary

In the present article, we attempted to replicate previous findings that hitting a target attenuates implicit motor learning. The results of the four experiments presented above suggest that implicit motor adaptation is modulated by task success as defined by the cursor hitting the target. The attenuation of learning driven by shifting the target to be concentric with the final cursor location (Target Jump paradigm) vastly exceeded the attenuation due to hitting a larger target that remained stationary (Target Size paradigm), indicating that these two different manipulations likely influence motor learning in different manners.

## ACKNOWLEDGEMENTS

This work was supported by a grant from the National Institutes of Health to OK (F32-NS122921) and the Blair Pyne Fund awarded to JAT.

